# Alzheimer’s subtypes A supervised, unsupervised, multimodal, multilayered embedded recursive (SUMMER) AI study

**DOI:** 10.1101/2025.05.09.653177

**Authors:** S. Kinreich, A. Bingly, G. Pandey

## Abstract

Since Alzheimer’s disease (AD) is a heterogeneous disease, different subtypes may have distinct biological, genetic, and clinical characteristics, requiring tailored interventions. While several proposed subtypes of AD exist, there is still no clear consensus on a definitive classification. By leveraging complementary AI approaches, including supervised and unsupervised learning, within a recursive pipeline (SUMMER) that integrates multimodal datasets encompassing MRI measurements, phenotypes, and genetic data, our goal was to generate robust scientific evidence for identifying AD subtypes. Data was downloaded from the Alzheimer’s Disease Neuroimaging Initiative (ADNI) database and included neuroimaging data (MRI), genetics (SNPs), clinical diagnosis, and demographics. 1133 European American participants’ images, aged 55-95, were included in this study. The analysis was multi-fold, where the first step involved applying an unsupervised application to a subset of the MRI sample (AD + cognitively normal (CN) aged matched groups, 100 men aged 68-85 years, and 76 women aged 68-85 years). The MRI brain gray matter was segmented into 44 regions of interest (ROIs) according to a standard atlas, and 618 features were extracted, including ROI voxel intensity measurements such as minimum, maximum, and histogram variables. Results identified a cluster of subtype AD men and a cluster of subtype AD women that were distinct from the rest of their respective samples. In the next step, the integrity of the identified subtype AD clusters was investigated using the XGBoost supervised machine learning application with genetic features (SNPs, N=36,724) and labels: the identified subtype AD cluster vs. the rest of the sample, stratified by sex. A significant AD subtype men model (accuracy=0.85, F1=0.72, AUC=0.83) and a significant women AD subtype model (accuracy=0.81, F1=0.81, AUC=0.81) were built, confirming the homogeneity of the isolated AD subtype clusters. Discriminative biomarkers were extracted from the significant models, including selected ROIs and SNPs. Finally, the subtype models were tested on an unseen subset of ADNI data. The genetic-based models identified clusters of AD subtype participants consisting of 34% of the men AD group and 47% of the women AD group. Phenotypic analysis indicates that lower body weight was associated with the women’s AD subtype. Complex diseases like AD demand a sophisticated, multimodal approach for precise diagnosis. Effectively identifying disease subtypes enhances the potential for personalized treatment, ultimately improving patient outcomes.

## Introduction

Cognitive decline in older adults is not attributed to a single neurological process but rather to several distinct subtypes^1^. One subtype is associated with natural aging^2^, while others are associated with neurological conditions such as Alzheimer’s disease (AD) and related dementia^3^. These diverse subtypes create ambiguity in diagnosing AD and understanding its origins, posing significant challenges for developing effective treatments and therapies^4^. It was estimated that in 2022, around 6.5 million Americans aged 65 and older were living with AD^5^. US forecasts indicate that the population of Americans with either Alzheimer’s dementia or mild cognitive impairment (MCI) will reach 15 million by 2060, up from 6.08 million in 2017^6^. These estimates highlight the critical importance of identifying risk and resilience biomarkers associated with each subtype’s underlying origins, enabling the personalization of future treatments and prevention strategies.

To this end, our work focused on phenotypic and biomarker differences between cognitive normal (CN) individuals and AD patients. We aim to build classifiers and predictive models to identify AD subtypes, distinguishing between AD and CN groups and subgroups within the groups to identify biomarkers associated with each.

Previous studies investigating AD subtypes suggested that based on pathologic measures of senile plaques, neurofibrillary tangles (NFTs, an abnormal accumulation of a protein called tau inside neurons)^7,8^, and cerebral amyloid angiopathy (CAA, a protein build-up on the walls of the arteries in the brain)^9^, 3 subtypes can be defined; typical AD, with balanced NFT counts in the hippocampus and association cortex^8^; limbic-predominant AD, with counts predominantly in the hippocampus^8^; and hippocampal-sparing AD, with counts predominantly in the association cortex^7,8^. Several neuroimaging studies have presented findings that support these subtypes by using visual rating scales of brain atrophy or automated methods for estimating regional brain volume^10,11^. A fourth subtype displaying minimal brain atrophy was also identified by neuroimaging studies^8^. Genetic studies have also explored the possibility of defining AD through distinct subtypes. For example, the MAPT H1H1 (microtubule-associated protein tau (MAPT) H1H1 genotype) was suggested to have a higher frequency in the limbic-predominant AD subtype compared with typical and hippocampal-sparing AD.^10,12^ However, another study reported that typical AD had the highest frequency of MAPT H1H1 carriers, followed by limbic-predominant and hippocampal-sparing AD^13^. The APOE gene E4 allele (APOE4) has been extensively studied and recognized as a major risk factor for late-onset AD^14^. Additionally, individuals who inherit two copies of the APOE4 variant face a substantially heightened risk of developing the condition^14^. With these encouraging results, the search for distinct AD subtypes continues, focusing on developing reliable and accurate diagnostic tools based on brain structure and function, genetics, and other phenotypes to identify AD subtypes effectively.

The complexity of identifying AD poses challenges when different subtypes are associated with different biological markers^15^. At the same time, biomarker analysis based solely on diagnosis and symptoms may be biased because symptoms are often similar between subtypes^15^.

Supervised machine learning algorithms may produce biased results due to misleading labeling that fails to account for the diversity of AD subtypes. Furthermore, the process of healthy aging can vary significantly in terms of brain anatomy and the timeline of changes, and this variation may also have an effect when interacting with disease subtypes^16^. To that end, unsupervised clustering can be effective, especially in the early stages when symptoms have not fully developed^17^. In the unsupervised approach, the software does not rely on prior knowledge that differentiates between groups, but rather looks for consistent patterns in the data to create clusters^18^. Clustering is a data-driven technique of grouping based on features that are similar to each other and separating them from those that are different^19^. Clusters are formed with features having minimum intra-cluster distance and maximum inter-cluster seperation^19^. An additional strength of the unsupervised learning approach is that it suffers less from data overfitting due to its embedded nature, where learning is done in an entirely blind manner^18^. Combining unsupervised and supervised methodologies may leverage the strengths of both approaches to effectively identify clusters within a dataset.

Moreover, our groundbreaking research^20^ and others^21^ have demonstrated that multimodal AI algorithms often outperform single-modality models in classifying complex diseases. By integrating diverse domains, such as environmental factors, genetics, and psychiatric diagnoses, our work leverages the strengths of each field while addressing their individual limitations, offering a more comprehensive framework for disease understanding and classification^20^. AD is a complex condition with potentially diverse origins, necessitating a multifaceted approach to identify and understand its various subtypes. In this study, we aimed to integrate multiple complementary approaches and datasets, including supervised, unsupervised, multimodal, multilayered embedded recursive (SUMMER) machine learning algorithms, brain structure, genetics, and phenotypic analysis, to uncover subtypes of AD.

To address this study’s aims, we used data from the Alzheimer’s Disease Neuroimaging Initiative (ADNI)^22,23^. The ADNI was launched in 2003 as a public-private partnership, led by Principal Investigator Michael W. Weiner, MD. The primary goal of ADNI has been to test whether serial MRI, positron emission tomography (PET), other biological markers, and clinical and neuropsychological assessment can be combined to measure the progression of MCI and early AD^23^. ADNI is a multisite, longitudinal study employing imaging, clinical, bio-specimen, and genetic biomarkers in CN as well as in individuals with early MCI and those who are diagnosed with AD^24^. ADNI follows individuals with and without AD and MCI, enabling a unique opportunity to assess an individual’s health status while incorporating age, sex, and ancestry differences into the analysis^22^.

To summarize, with ADNI’s rich genetic and phenotypic data from multiple domains (e.g., clinical, neuroimaging (MRI), psychosocial, psychiatric, and demographic), we applied brain-based unsupervised algorithms that have the potential to identify biodiversity patterns related to dementia subgroups that would otherwise remain hidden. We utilized the ADNI’s individual genetics information to further confirm the AD subtype’s robustness initially found by the clustering of the unsupervised machine learning algorithm. Our multilayered analysis continued and validated the results with recursive analysis, where the AD subtype finding was validated with unseen ADNI data.

## Method

We present here an assembled pipeline constructed from compatible machine learning applications (unsupervised and supervised) that were applied to various modalities (neuroimaging, genetics, and phenotypes) in order to identify and establish distinctive AD subtypes. All calculations were done using Python packages and FreeSurfer (version 4, http://surfer.nmr.mgh.harvard.edu, Boston, MA)^25^.

### ADNI Sample

The MRI data acquired from AD and CN participants were downloaded from the ADNI database (https://adni.loni.usc.edu/)^26^. Only European American (EA) participants were included in the analysis due to the low number of other ancestries in the ADNI dataset, and only participants with genetic information, resulting in an initial dataset that included 1,113 brain images of participants between the ages of 55.1 and 95.8 years. Initial data included multiple visits by each participant (588 men’s brain images and 525 women’s brain images). Details of the ADNI design, participant recruitment, clinical testing, and additional methods have been published previously^22,27^ and are available at www.adni-info.org.

### MRI scans

Raw ADNI 1.5 T and 3.0 T MRI scans were downloaded from the ADNI public database (www.loni.ucla.edu/ADNI). The raw DICOM files were transformed into NIFTI format^25^.

### MRI preprocessing region segmentation

The MRI data was preprocessed using standard procedures that included 1. skull-stripping using the Brain Extraction Tool (BET)^28^ (Fig 1A). 2. Brain normalization to standardize all brains to the same scale. 3. Brain registration to the standard template (MNI152_T1_1mm_brain) using the Python package of ANTS^29^. This process harmonized the data into a voxel resolution of 1mm^3^. 4. Brain segmentation according to the structural standard brain template, defining 44 ROIs for each brain^30^. 5. The ROIs masks were created using The FSLView^**31**^ with the same brain template (MNI152_T1_1mm_brain) and atlas (Harvard-Oxford cortical and subcortical structural atlases)^30^ (Fig 1B). ROIs included cortical and subcortical structural areas according to the atlas^30^ (full list in the supplementary file). Only selected available ROIs were extracted separately from the left and right hemispheres (amygdala, thalamus, putamen, hippocampus, caudate, and nucleus accumbens), while the rest were averaged across the hemispheres.

**Figure 1.**
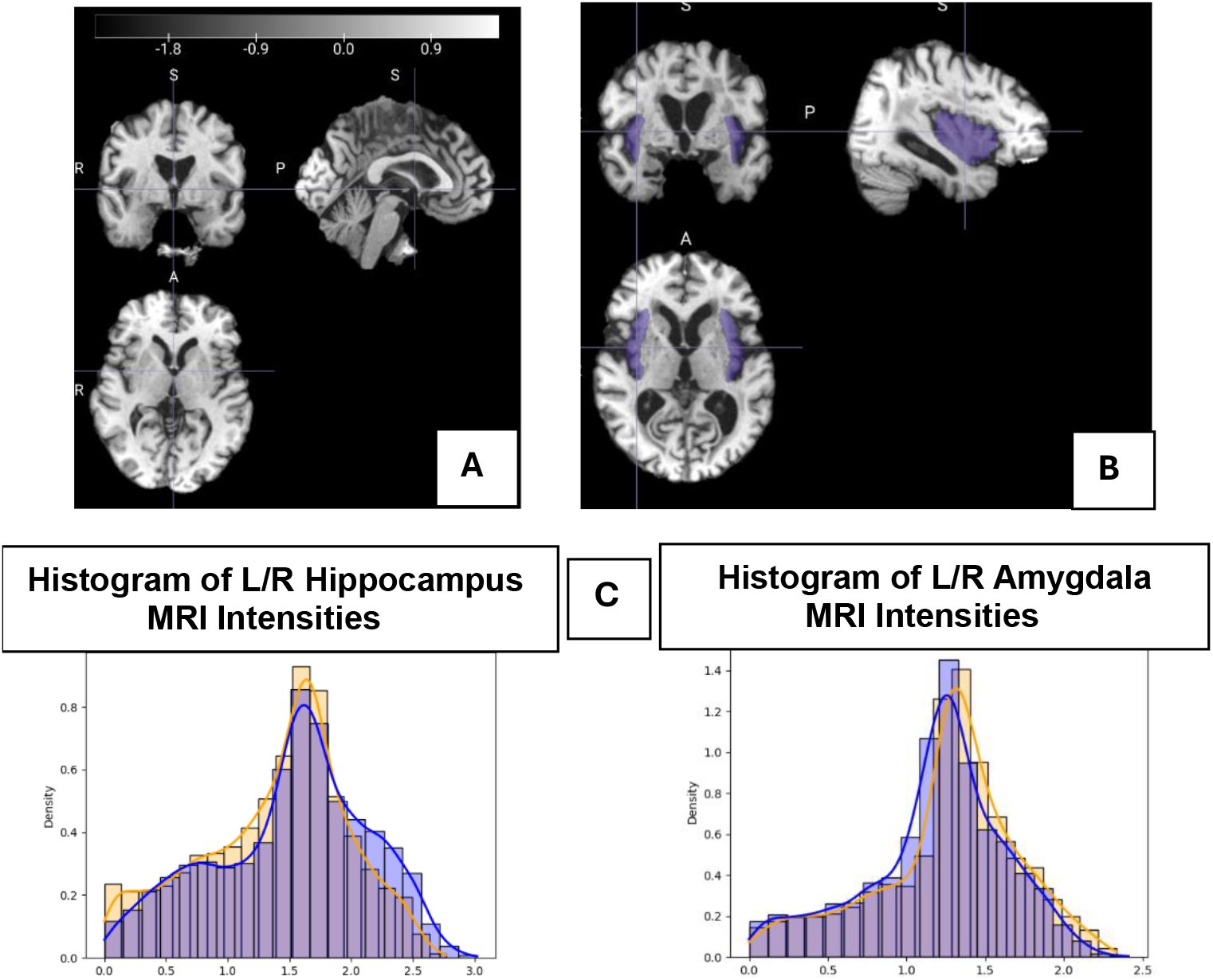
Brain preprocessing and ROI extraction. A. The brain image was stripped from the scalp, registered to a standart template, and normalized. B. 44 ROIs intensities values were extracted from each brain. Here is an illustration of the extracted insula ROI colored in purple. C. Histogram of intensities of the contralateral Hippocampus (on the left) and Amygdala (on the right). Right ROI, Orange. Left ROI, blue and shared area; purple.

### MRI ROI measurements

The most straightforward way to examine brain anatomy is through visual inspection of MRI scans^32,33^. However, this method is time-consuming, requires substantial operator training, and suffers from inconsistent tracing protocols across laboratories^32^. Moreover, research suggests that visual inspection may be subject to perceptual bias^34^. To address these challenges, automated software tools that analyze MRI intensities have been developed to assist in identifying tumors^35^, delineating ROI volumes^36^, and quantifying other brain variables^37^. MRI voxel intensities are rarely used directly as features in neuroimaging research, with only a small subset of Alzheimer’s disease studies employing voxel-based intensity analyses^38-40^.

Compared to functional MRI (fMRI), voxel intensity analysis does not require complex task-based experiments and can be applied to any standard structural MRI scan^31^. Following the preprocessing and harmonization/normalization steps that allowed the MRI scans to be on the same scale, the following measurements of the MRI voxel values were calculated: minimum, maximum, and standard deviation. Additional measurements included values within the bottom 10^th^, 20^th^, and 25^th^ percentiles, as well as the top 10^th^, 20^th^, and 25^th^ percentiles. These values were calculated to reduce the possibility that ROI’s intensity outliers will bias the results^41^.

Additionally, histograms were calculated for each ROI’s (Fig 1C) voxel values, and the histogram entropy was added to the features (see Table 1 in supplementary file for feature list). Next, for selected ROIs (amygdala, thalamus, putamen, hippocampus, caudate, and nucleus accumbens), the difference between the contralateral histogram ROIs was calculated.

**Table 1.** The Unsupervised Analysis Sample.

| **Table 1. The Unsupervised Analysis Sample** |  |  |  |  |  |
| --- | --- | --- | --- | --- | --- |
| | # CN images | # AD images | Mean Age CN ± SD | Mean Age AD ± SD | P value |
| Men | 50 | 50 | 78.12±4.37 | 77.83±4.85 | 0.74 |
| Women | 38 | 38 | 76.53±4.18 | 76.52±4.19 | 0.99 |
| CN: Cognitively normal; AD: Alzheimer's Disease; SD: standard deviation |  |  |  |  |  |

Functional and morphological differences between hemispheres have previously been suggested in the study of various brain disorders^42^. Contralateral measurements included the Euclidean distance, which is the difference between the entropy of histograms, the percentage of mutual area (vs. both histograms), the percentage of mutual areas (vs. the right), and the percentage of mutual areas (vs. the left) (Fig 1C). Overall, the analysis included 618 features for each ROI. The features were randomly divided into 36 groups of nine to maximize their individual contributions^43^ and the ML algorithm iterated over these groups. A cluster was defined as visibly isolated from the other participants and included more than 85 percent of AD participants. This threshold was chosen arbitrarily to ensure the inclusion of the majority of the AD participants in the cluster.

### Genetics

Raw genetic information was downloaded from the ADNI public database (www.loni.ucla.edu/ADNI). The Human 610-Quad BeadChip (Illumina, Inc., San Diego, CA) was used to genotype all ADNI participants in the database, which resulted in a set of 620,901 SNPs^44^ for each. The specific methods related to ADNI genotyping can be found on the ADNI website (https://adni.loni.usc.edu/data-samples/adni-data/genetics-related-omics).

To further study the homogeneity of the MRI-based isolated cluster as an AD subgroup, we investigated the genetic profile of the cluster compared to the rest of the group. Supervised machine learning analysis (XGBoost) was applied with SNPs as features and with the isolated group and the rest of the group as labels. The SNPs were first normalized, and then SNPs with missing values (more than 5% for men, 10% for women) were discarded. Next, we used the least absolute shrinkage and selection (LASSO) method for feature selection^45^. The sparsity property of LASSO (i.e., most coefficient estimates being exactly zero) makes it attractive for feature selection, as it reduces the estimated variance while providing a more interpretable final model^46^. Its applications to genomic data^47^ and behavioural markers^48^ have shown that selecting a small number of representative features can achieve more accurate classification than all the features. We first determined the LASSO regularization parameter using a 10-fold cross-validation (CV) procedure with labels “isolated cluster” and “rest of the group” as the response variable. All the features with a non-zero coefficient were retained for subsequent analyses.

### Demographic and clinical data

Demographic and diagnosis information for all analyzed visits was downloaded from the ADNI clinical data repository (https://www.loni.ucla.edu/ADNI/Data/ADCS_Download.jsp).

### SUMMER Analysis Pipeline

The outline of the AD subtype pipeline (including supervised AI, unsupervised AI, multimodal, multilayered embedded recursive approaches (SUMMER) is presented in Figure 2.

**Figure 2.**
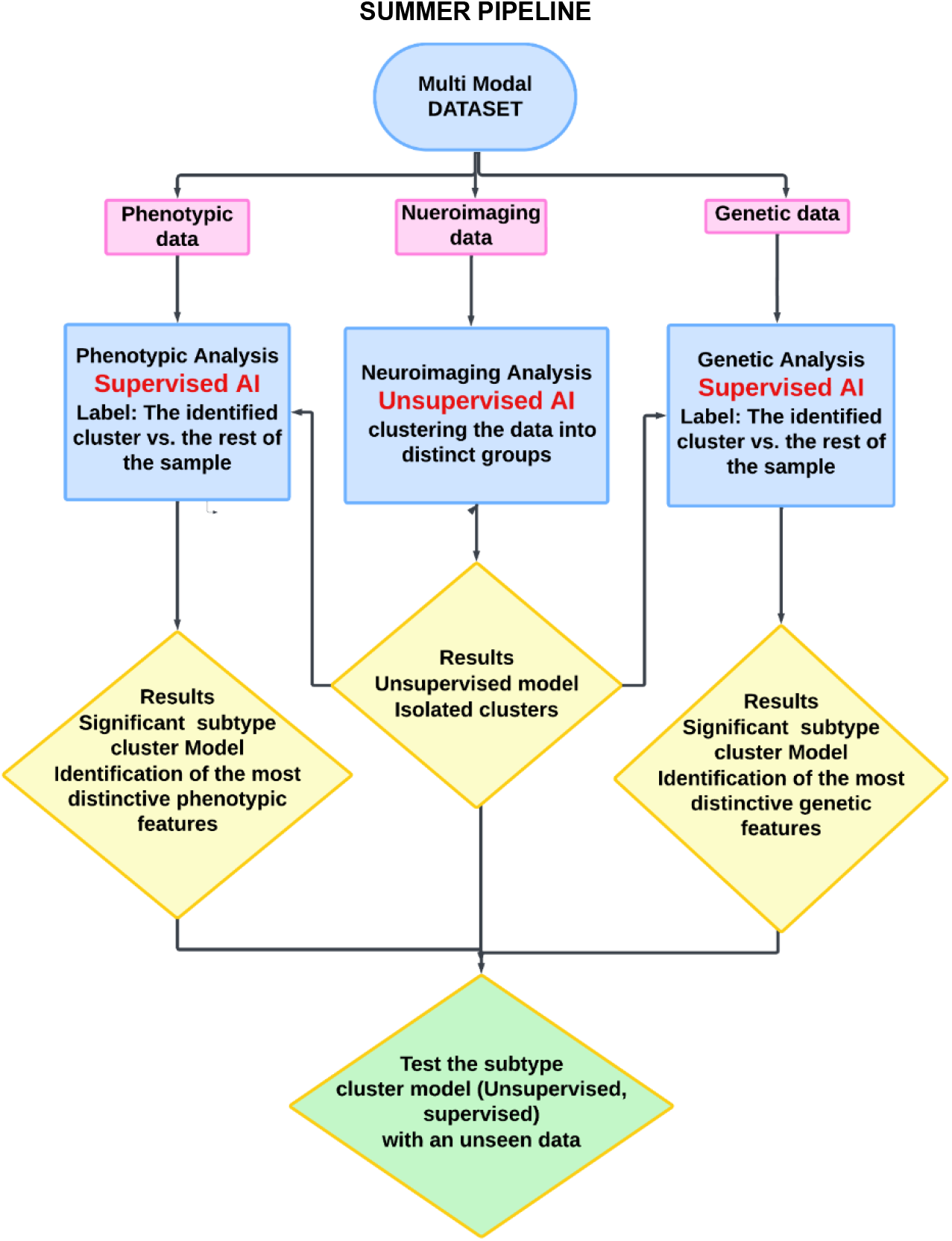
SUMMER pipeline

Following the SUMMER pipeline, we conducted a series of AI analyses, each incorporating different samples and modalities, with each analysis building upon the results of the previous one. We first identified the AD clusters using an exploratory unsupervised approach with MRI data. To ensure the integrity of the clusters, we then validated them using a supervised approach incorporating genetic data.

Finally, we tested the models on an unseen ADNI data subset to ensure robustness and generalizability. Below is the breakdown of the steps.

### 1. Unsupervised machine learning clustering with structural (MRI) features

AI unsupervised learning applications in machine learning allow the interpretation of data without predefined labels^19^. The applications rely on algorithms that identify patterns, structures, and relationships in the data^18^. Our sample included ADNI Age-matched AD and CN participants, stratified by sex (men, women), ranging from 68 to 85 years old^49^ (See Table 1 for participant information). The groups (AD, CN) included two visits from each participant to increase the sample size. The age range constraint was applied based on previous findings, confirming age’s significant impact on brain structure and function^50^. Similarly, previous findings on sex differences in brain function were the reason for stratified analysis by sex^51^. A total of 618 features were computed per ROI (see Methods). Given 44 ROIs per participant, this resulted in 27,192 features per participant. The dimensionality of the features was reduced by performing a t-test on all features, comparing AD to CN to focus on AD-related characteristics. Only features that met the significance threshold (p < 0.05) were included in the unsupervised analysis, resulting in a final set of 50 MRI features. We applied the Uniform Manifold Approximation and Projection for Dimension Reduction (UMAP)^52^ unsupervised algorithm with the MRI features. UMAP^53^ is a software tool for unsupervised dimensionality reduction, which is pivotal for visualizing and clustering unannotated high-dimensional data^53^. Dimensionality reduction is aimed at transforming the data into a relevant and reduced dimensional space by discovering intrinsic data structure and limiting redundancy. It allows extracting useful variables while discarding noise and correlated features^53^. UMAP is a manifold learning technique based on Riemannian geometry and algebraic topology. UMAP was used with the participant’s MRI ROI values to capture the underlying patterns. The ROIs’ features that contributed most significantly to cluster formation were identified by performing a t-test between the cluster and the rest of the data for each significant model (men and women). A false discovery rate (FDR) was applied to correct the results for multiple comparisons^54^.

### 2. Supervised machine learning with genetic data

Here we aimed to reinforce the AD cluster and its integrity further by investigating its genetic homogeneity compared to the rest of the sample, which includes both CN and AD participants. The sample included the same cohort from step 1, i.e., age-matched AD and CN participants, stratified by sex, ranging from 68 to 85 years old, was included in this analysis.

Features included ADNI’s genetic data (SNPs)^55^ Following the LASSO application, all the features with non-zero coefficients were retained for subsequent analyses, resulting in ∼20 SNPs for men and ∼20 SNPs for women.

The Supervised eXtreme Gradient Boosting (XGBoost)^56^ was applied with the isolated cluster vs. the rest of the group as labels. XGBoost is an open-source machine learning library that uses gradient-boosted decision trees^56^. It is known for being fast, efficient, and able to handle large and small datasets^57^. The reduced set of features identified was fed to the XGBoost algorithm to classify the isolated cluster.

### 3. Testing the genetic-based model with ADNI unseen data

In this step, the cluster model built by the genetic-based supervised algorithm was tested on an independent dataset comprising all ADNI participants who were not included in the previous analysis and did not serve as the model’s foundation (N men =365, N women=438) with age range of 55.1 and 95.8. The sex-specific models were applied with their corresponding sample and features (SNPs)(more information about the sample is given in the supplementary file). The model was expected to identify a subtype of AD that would include AD patients.

### 4. Phenotype analysis

The subtype clusters were also tested for phenotypic (body weight) association with AD. Evidence suggests that body weight can impact the risk and progression of AD. Being overweight or obese in midlife is linked to a higher risk of developing AD later^58^. Excess weight can contribute to conditions like high blood pressure, diabetes, and inflammation, all of which are associated with cognitive decline^59^. On the other hand, People with AD often experience unintentional weight loss, especially in the later stages^60^. This can be due to decreased appetite, difficulty eating, changes in metabolism, or trouble recognizing food^60^. To test the relationship between AD and body weight, a t-test was conducted between the recorded weight data of the participants in the cluster and the weight data of the rest of the sample.

## Results

### 1. AI Unsupervised algorithm

We applied UMAP^52^ unsupervised algorithm with the MRI features. All results (distribution of the participants’ UMAP values in 2d space) from the unsupervised analysis can be found in the supplementary file. Results for the women group revealed an isolated cluster composed mainly of AD participants (86% AD, Figure 3A). Results for the men group revealed an isolated cluster composed mainly of AD participants (92.3% AD, Figure 4A). The features that most strongly contributed to the formation of the distinct cluster of men and women AD subtypes were found to be primarily associated with brain networks known to underlie AD-related memory function, including regions such as the hippocampus and parahippocampus. The fusiform gyrus, traditionally associated with face recognition processes, was also found to be linked to both sexes AD subtypes. The women AD subtype was found to be associated with the thalamus and the caudate and men-specific AD subtype was associated with the parietal operculum (Figure 3B (women) and 4B (men)). Overall, all MRI average values were lower for the AD cluster subtype and higher for the SD values and entropy values for men and women. Table 2 in the supplementary file presents the isolated cluster vs. the rest of the data mean values and p-values, and Table 3 in the supplementary file presents the mean values and the p-values of all brain areas associated with the clusters.

**Table 2.** XGBoost Supervised AI Model Cluster Accuracy.

| Table 2. XGBoost Supervised AI Model Cluster Accuracy |  |  |  |  |  |
| --- | --- | --- | --- | --- | --- |
| Sex | Precision | Recall | Accuracy | F1 | AUC |
| Men | 0.66 | 0.8 | 0.85 | 0.72 | 0.83 |
| Women | 0.75 | 0.85 | 0.812 | 0.81 | 0.81 |

**Table 3:** AD subtype of women and men, selected features.

| Table 3: AD subtype of women and men, selected features |  |  |
| --- | --- | --- |
|  | Women AD subtype | Men AD subtype |
| **ROI** | top 25<sup>th</sup> Parahippocampal Gyrus | top 25<sup>th</sup> Supracalcarine Cortex |
|  | Entropy Parahippocampal Gyrus | STD Occipital Fusiform |
|  | top 20<sup>th</sup> Temporal Fusiform | top 10<sup>th</sup> Lingual Gyrus |
|  | bottom 20<sup>th</sup> Heschl Gyrus | top 20<sup>th</sup> Parietal Operculum |
|  | Mean Right Caudate | STD Right Hippocampus |
|  | Entropy right Hippocampus |  |
|  | Entropy Superior Temporal |  |
| **SNPs** | rs11689050 | rs7722673 |
|  | rs1874161 | rs1538608 |
|  | rs10216685 | rs2396274 |
|  | rs6775796 | rs1122269 |
|  | rs7306261 | rs17131515 |
|  | rs2139011 | rs10050582 |
|  | rs10069215 | rs17513184 |
|  | rs2422973 | rs12130821 |
|  | rs17456902 | rs9425322 |
| **Body weight** | Lower body weight |  |

**Figure 3.**
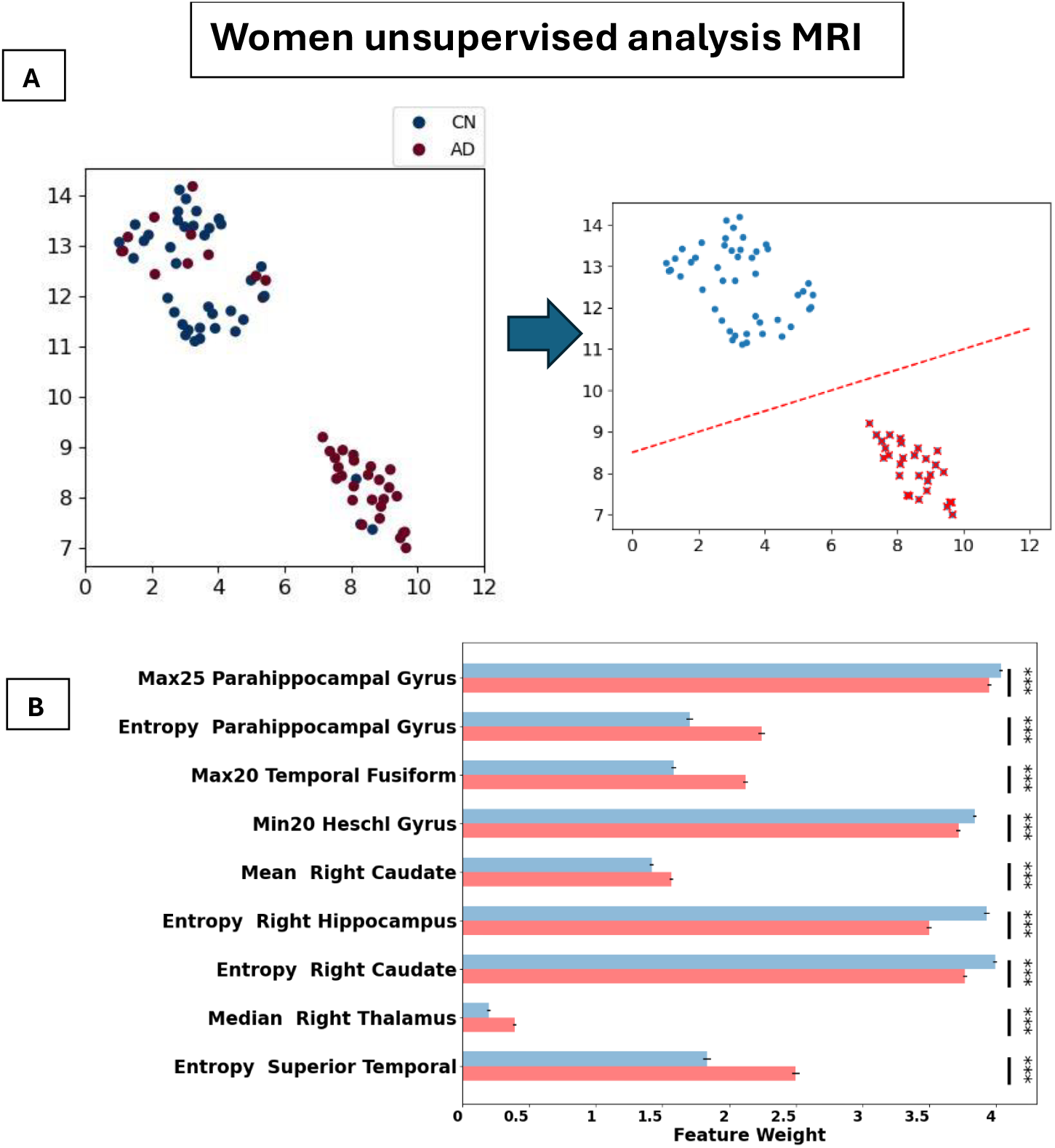
The women’s unsupervised clustering of the MRI measurements revealed an isolated AD cluster. A. It is evident that the majority of the isolated cluster includes AD participants. On the right, the isolated cluster is colored in red, and the rest of the sample is colored in blue. B. The bar graph presents ROIs that contributed the most to the creation of the cluster. Max25-top 25^th^ percentile, Max20-top 20^th^ percentile, Min20-bottom 20^th^ percentile.

**Figure 4.**
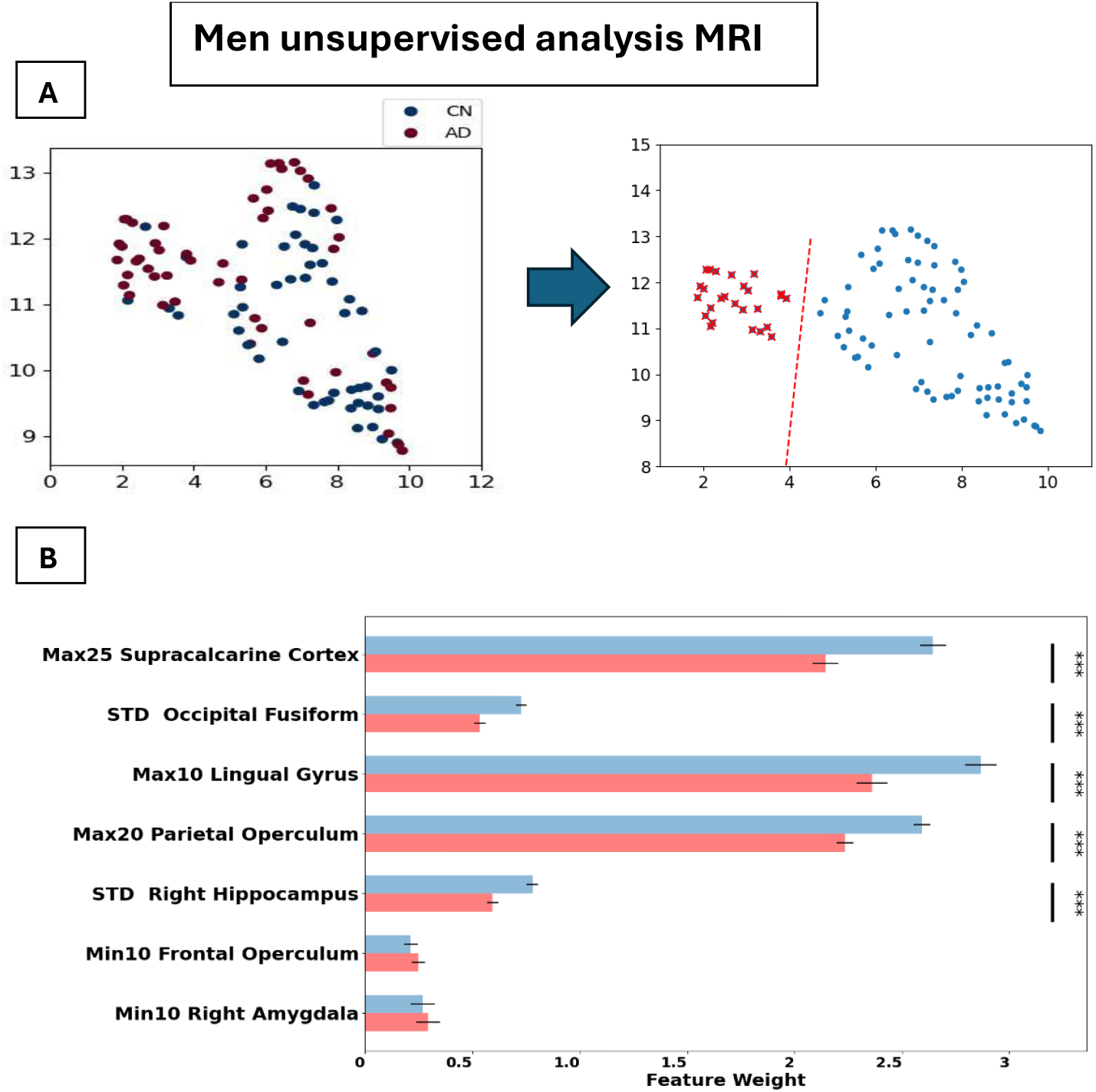
The men’s unsupervised clustering of the MRI measurements revealed an isolated AD cluster. A. It is evident that the majority of the isolated cluster includes AD participants. On the right, the isolated cluster is colored in red, and the rest of the sample is colored in blue. B. The bar graph presents the ROIs that contributed the most to the creation of the cluster. Max25-top 25^th^ percentile, Max20-top 20^th^ percentile, Max10-top 10^th^ percentile, Min10-bottom 10^th^ percentile. STD − Standard deviation

### 2. Supervised machine learning with genetic data

Significant models were created for both men (accuracy=0.85) and women (accuracy=0.81). Table 2 presents model information, including precision, recall, F1 score, and area under the curve (AUC), confirming the genetic heterogeneity of the clusters. The most important features contributing to the formation of the clusters include SNPs such as; (men) rs7722673 (gene WWC1), rs1538608, rs2396274, rs1122269, and rs1122269 (gene CDH4), and (women) rs11689050 (gene LINC01808) rs1874161, rs10216685, rs6775796, rs2422973 (gene RASSF2), rs12130821 (gene RGS5), rs9425322 (gene RGL1), rs17794576 (gene NCAM2), rs7306261 (Figure 5). A full, detailed SNPs summary is presented in Supplementary Table 4.

**Figure 5.**
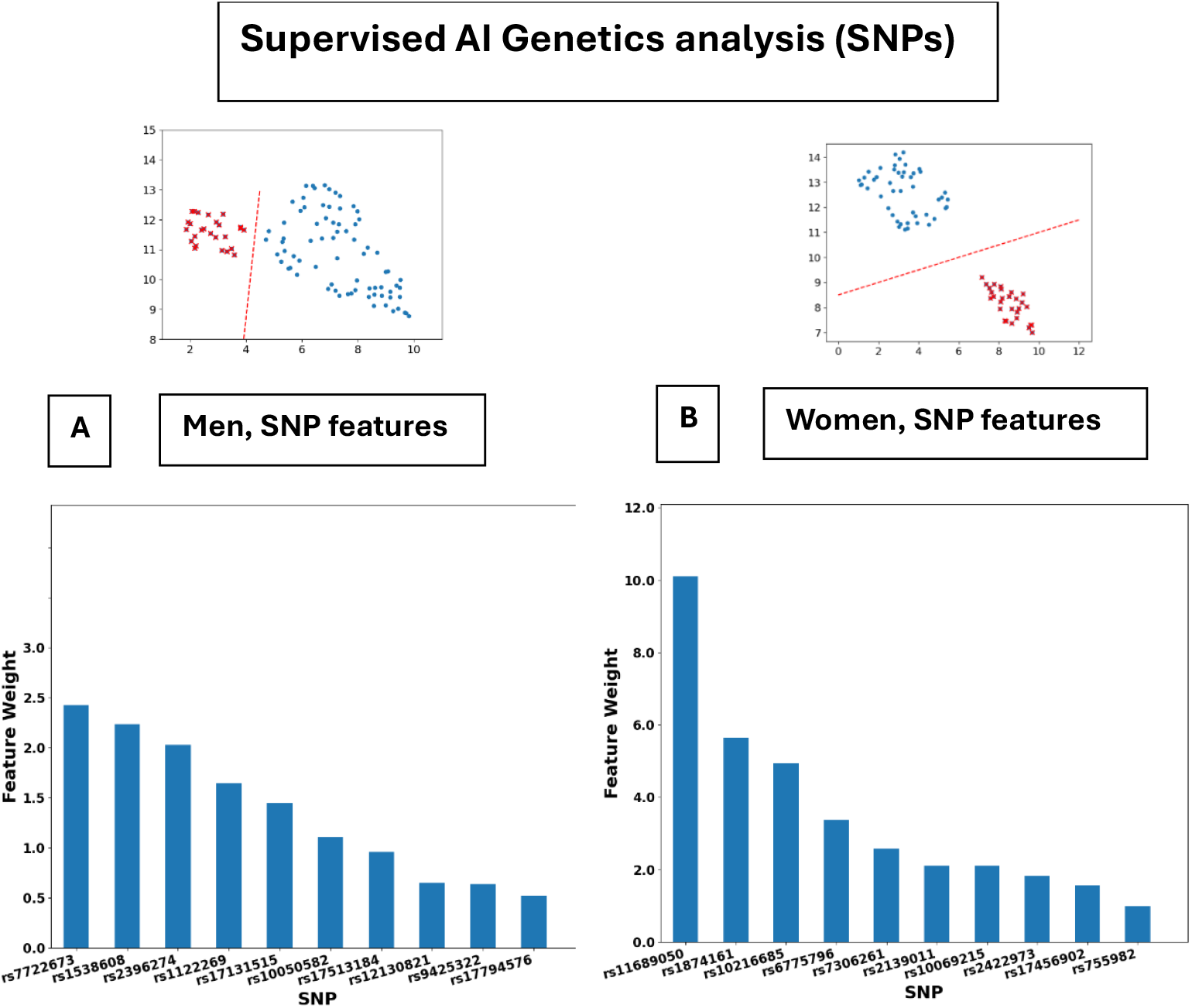
XGBoost Supervised Al Genetics Analysis (SNPs). The XGBoost analysis labels were the cluster (colored in red) vs. the rest of the sample (colored in blue). A. SNPs that are the most significant contribution to the XGBoost model for men (A) and women in (B)

### 3. Testing the genetic-based model with ADNI unseen data

The genetic-based models (men and women) were tested with unseen data. Figures 6A and 6B show the sample percentage of AD subtype clusters that were identified by the men and women models, respectively. Results show that 47% of the women subtype cluster comprised AD participants, while in the rest of the data included 10% AD participants (Figure 6). 30% of the men cluster were AD, while 15% were AD in the rest of the sample (Figure 6 and see Table 5 in supplementary file).

**Figure 6.**
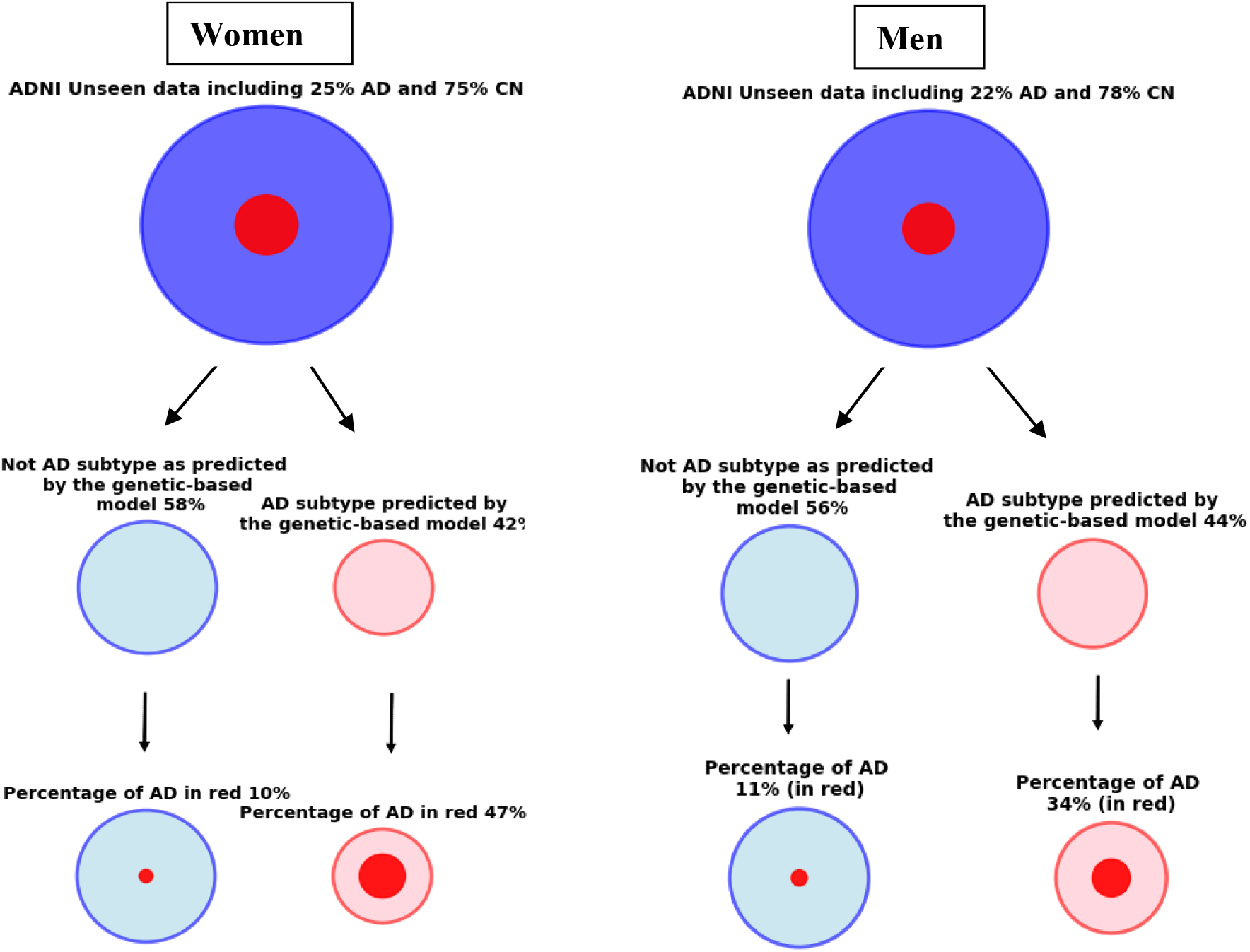
Genetics-based AD subtype model applied with ADNI unseen dataset. For both sexes, the models identified a cluster that included AD patients.

### 4. Phenotypic analysis

Results show an association between the participant’s weight and the women AD subtype cluster, p = 0.004 (AD, mean=63.29±9.85, CN, mean=71.31±13.65). No association was found between men AD subtype cluster and body weight, p == 0.6 (AD, mean = 83.54±9.82, CN, mean=82.14±13.13 (Table 3. for full subtype biomarkers and phenotypic description)

## Discussion

Diagnosing AD subtypes, especially in the early stages of the disease, holds enormous potential for research on drug interventions and treatments. By mining high-content data sets using an exploratory, unsupervised data analytics approach, we were able to distinguish AD subgroups and identify their respective biomarkers. The world of medicine strives for more personalized medicine, identifying the needs of the individual over the “average” treatment in order to improve treatment results. In our innovative SUMMER model analysis, we have assembled different types of AI and multimodality approaches, incorporating multilayered and recursive steps enabled by the multimodal ADNI dataset. We identified distinctive subtype clusters of men AD and women AD by an unsupervised analysis applied to MRI intensity measurements. The most discriminative features in the AD subtype models include MRI measurements from ROIs, such as the hippocampus and the fusiform gyrus. The AD subtype cluster homogeneity was confirmed with a significant genetics-based (SNPs) XGBoost supervised model. AD subtype model-specific SNPs were identified for each of the sexes. Phenotypic analysis of body weight found that the women AD subtype has lower body weight than the rest of the sample. This association was not found for the men’s AD subtype. The significant genetic models were tested with ADNI unseen data identifying a men AD subtype cluster and women AD subtype cluster in this dataset. The results highlight the superiority of multimodality biomarkers in identifying subtypes of complex diseases such as AD.

### Multi-modality features and AD subtypes

In this study, we demonstrated that utilizing multiple AI approaches and modalities allowed the identification of homogeneous clusters. Specifically, the unsupervised approach successfully classified AD subtypes that would have otherwise remained hidden within the averaged results. Our previous groundbreaking study on predicting the development of Alcohol Use Disorder^20,61^, along with others^21^, demonstrated that models integrating multimodal data, such as neuroimaging biomarkers and genetic information, outperform those relying on a single modality^20^. These findings align with the understanding that complex diseases are influenced by multiple factors, including genetic and environmental components^62^. Our results indicate that distinct AD subtypes can be effectively identified using a series of AI algorithms applied sequentially, and in a recursive manner across multiple datasets. An alternative AI approach explored in recent studies, also utilizing multi-modal data, is implemented as a single-layer model rather than a sequential one. In the one-layer approach, multiple features from different modalities are reduced to display an efficient set of features to predict/classify the labels. This reduction due to collinearity may cause a loss of information. For example, when building a model that is composed of multiple modalities, such as phenotypes, brain functions, and genetics, the trained model may be based on only some of the modalities. Additionally, certain modalities are naturally more sensitive to factors such as age and sex. By adopting a sequential approach, we can more effectively balance and manage the trade-offs imposed by these constraints. For example, in our study, we applied unsupervised algorithms to neuroimaging data across a restricted age range, which was necessary due to the significant impact of age on brain structure.

### AD subtype-related brain features

AI unsupervised analysis revealed a set of ROIs associated with AD subtypes in both sexes. The identification of subtype-specific features, including brain ROI measures and genetic factors, can provide valuable insights into the onset and progression of AD. Our findings show that men and women AD subtypes have been found to be associated with known AD-memory-related brain networks, such as the hippocampus and parahippocampal gyrus, located in the medial temporal lobe^63^. These brain structures play a crucial role in forming and consolidating new explicit (declarative) memories (e.g., facts and events)^64^. The hippocampus is consistently implicated in AD-related brain structure, function, and behavioral studies, showing reduced volume^65^, rapid tissue loss^66^ and functional disconnection with other brain regions^67^. The fusiform gyrus, traditionally associated with face recognition processes, was also found to be linked to AD subtypes in both sexes. The fusiform gyrus is located below the lingual and parahippocampal gyri, both also associated with the AD subtypes, and together, they have been highlighted in imaging studies, demonstrating a strong link to memory impairment^68,69^. Fusiform may shrink in people who develop AD^70^ and genes within the fusiform gyrus may be involved in AD onset and progression^71^. The fusiform gyrus was also implicated in mild cognitive impairment and loss of facial recognition^72^, both of which are associated with the development of AD. The women AD subtype was found to be associated with the thalamus and the caudate. The thalamus functions as a relay center, transmitting sensory and motor signals to the cerebral cortex and the caudate nucleus^73^, a crucial part of the basal ganglia, involved in processing spatial information and regulating motor behavior^73^. Both brain structures were previously implicated in AD, showing volume reduction^74^. Furthermore, a recent imaging study revealed that the amyloid imaging marker AV-45, a PET imaging agent used to visualize beta-amyloid plaques in the brain, is elevated in the caudate nucleus and putamen of individuals experiencing rapid cognitive decline^75^. More women-specific AD subtype brain regions include the Heschl’s gyrus. Heschel’s gyrus processes the frequency, duration, and intensity of sounds and changes in hearing, especially speech perception in noise, all of which may be a preclinical marker of neurodegeneration.^76^ More so, this brain structure was found to be associated with AD in a volumetric MRI study^77^. Men-specific AD subtype was associated with the parietal operculum, a parietal lobe region. The parietal lobe was implicated in AD, and its dysfunction contributes to memory, language, and visuospatial deficits, particularly in the early stages^78^. In a transcranial magnetic stimulation(TMS) study, the parietal operculum specifically has been found to be involved in proficient memory-guided haptic grasping^79^. This finding and others^80^ led to the suggestion that the parietal operculum acts as a “hub,” where several sensory-motor streams originating from different cerebral areas converge^80^. Our results add to these accumulating AD-related findings, suggesting that this group of ROIs characterizes the specific AD subtype revealed in the unsupervised analysis.

### AD subtype-related genetics

The results show that the AD subtype clusters were genetically identified by a collection of SNPs, confirming the subtype’s generalizability. Women-specific SNPs included a locus on chromosome 2, gene Long Intergenic Non-Protein Coding RNA 1808 (LINC01808) (rs11689050, weight ranking 1). In GWAS analyses, LINC01808 has been found to be associated with phenotypes such as height^81^, pulse pressure measurement^82^, ascending thoracic aortic diameter^83^, and directly relevant to the current study, AD (MTAG)^84^, and PHF-tau measurement^85^. PHF-tau, or paired helical filament tau, is a biomarker for AD, and its measurement involves detecting hyperphosphorylated tau protein, which forms neurofibrillary tangles^86^. The AD women subtype was also associated with a locus on chromosome 20, gene RASSF2 (rs2422973, weight ranking 8). This gene is associated with DNA methylation variation (age effect)^87^ and encodes a protein that contains a Ras association domain (rheumatoid arthritis (RA) protein)^88^, which plays a crucial role in signal transduction pathways (EGLN1/RASSF2 protein level ratio)^89^. The men AD subtype was associated with a locus on chromosome 5, gene WWC1 (rs7722673, weight ranking 1). A GWAS study found that WWC1 gene is associated with protein KIBRA levels^90^. KIBRA is a protein involved in learning and memory, and its levels are associated with cognitive impairment and tau pathology in conditions like AD^90^. WWC1 was also found to be associated with brain morphology in a schizophrenia study^91^ and educational attainment^92^. Another locus was found on chromosome 1, gene CDH4 (rs1122269, weight ranking 5). CDH4 encodes a protein called retinal cadherin (R-cadherin) which is a cell adhesion protein that plays a crucial role in maintaining the structural integrity of tissues and organs, particularly in the retina and brain^93^. Studies show an association of this gene with educational attainment^94^. Both men and women subtype clusters found genetic association with smoking initiation (CDH4 and LINC01808 genes, respectively).^95^ A locus on chromosome 1, RGS5 gene (rs12130821, weight ranking 8) has been associated with height in a diverse ancestry GWAS^81^. In the same GWAS, gene RGL1 (rs9425322, weight ranking 9) was found to be associated with height. In a GWAS of both sexes, RGS5 was associated with testosterone in the men group^96^. In a VA Million Veteran Program GWAS, RGS5 was found to be associated with lipoprotein disorders (PheCode 277.51)^97^. A locus was found on chromosome 21, gene NCAM2 (rs17794576, weight ranking 10). The protein encoded by this gene belongs to the immunoglobulin superfamily. It is a type I membrane protein and may function in selective fasciculation and zone-to-zone projection of the primary olfactory axons. Even though the revealed men/women AD subtype clusters do not share genes, they share several phenotypes such as height, smoking initiation, and educational attainment. Our data-driven subtype cluster analysis revealed a select set of genes previously implicated in AD and dementia. The findings specifically suggest that these genes are linked to a distinct subtype of AD, significantly distinguishing it from other types of AD.

### AD subtype and phenotypes − body weight

An association with weight was observed in the identified women AD subtype but not in the men one. The revealed women’s AD subtype cluster was associated with lower weight compared to the rest of the ADNI sample. These results confirm previous studies showing that clinically important weight loss occurs more frequently in patients with AD than in cognitively normal control subjects^60^. Along the same lines, a meta-analysis that included 19 studies with 589,649 participants who were followed for up to 42 years concluded that, compared to a healthy weight, being underweight in midlife was associated with a 39% higher risk of dementia^98^. More broadly, there is evidence of a complex relationship between weight and AD/dementia^99^. For example, obesity or being overweight during middle age (roughly 35-65) can increase the risk of developing dementia later in life^100^, while late-life obesity might be associated with reduced dementia risk^101^. Particularly in women, one of the factors that has been suggested to influence the association between body weight and dementia/AD is hormonal changes that occur during menopause; specifically, estrogen depletion has been found to be associated with obesity and the risk of AD^102^. Several factors were suggested for weight loss in people suffering from dementia/AD^103^, such as reduced energy intake owing to their decreased mental status^104^, sensory changes such as deterioration of the olfactory bulb and/or diminished gustatory perception as a result of cholinergic deficits^105^, or dysphagia, or difficulties swallowing, may reduce energy intake^106^. AD-related increase^107^ or decrease^108^ body weight was also found to be associated with hypermetabolism, defined as an elevated basal metabolic rate of >10% or neuroendocrine dysregulation^109^. Our theoretical model proposes that what was once considered a single entity (AD diagnosis) is actually a collection of distinct subtypes, each defined by unique physiological and environmental variables. Our findings indicate that the identified female AD subtype is associated with lower body weight, along with distinct MRI biomarkers, and specific genetic variants (SNPs/genes). The revealed AD men subtype was not found to be associated with weight changes. Achieving a deeper understanding of each AD subtype requires integrating information from multiple modalities to distinguish between them effectively.

The SUMMER pipeline enables modular integration of different modalities and algorithms, optimizing the identification of AD subtypes. Relying on a single algorithm or combining all methods into one comprehensive model may obscure important distinctions, such as between different age groups or due to comparability that may eliminate key variables from the model. Integrating unsupervised algorithms into the analysis protocol is particularly beneficial for subtyping research, as these methods can explore large datasets in a blinded manner, independent of clinical diagnoses or self-reported information, and identify homogeneous clusters that help distinguish between different types of disorders. In this study, we identified distinct subtypes of men and women AD, each characterized by unique SNPs profiles and phenotypic characteristics. Recognition of these subtypes opens the door to earlier detection and the potential for personalized treatment strategies.

### Limitations and future directions

We present a pipeline applied to the ADNI dataset, yielding promising results. However, the identified AD subtype may be influenced by the small sample size. Future research with larger datasets should reevaluate the generalizability of these findings and validate them. Additionally, further studies should be conducted to identify distinct subtypes that can be clearly differentiated from the ones we uncovered. These subtypes are likely to be associated with a different set of genes and a different set of brain structures. Furthermore, a comprehensive investigation is required to clarify the relationship between AD subtypes and different phenotypes, which will allow for a more precise definition of each subtype. One key research decision was to apply the pipeline to a mixed dataset comprising both CN and AD participants rather than exclusively to AD cases. While both approaches are valid, we chose the mixed dataset to demonstrate the model’s applicability to individuals before establishing an AD diagnosis. Future research applying the SUMMER Pipeline to an AD-only sample would deepen understanding of AD subtypes.

## Acknowledgments

Sivan Kinreich discloses support for the research from NIAAA AA029448.

## Notes

https://adni.loni.usc.edu/data-samples/adni-data/

### Competing Interest Statement

The authors have declared no competing interest.

### Summary of Updates

The text was updated to correct several statements. First, it was clarified that SNP rs1122269 is located in an intron of CDH4 (both in the main text and in Supplementary Table 4). Additionally, in Supplementary Table 2, the column headers "Mean CN" and "Mean AD" were corrected to accurately match their respective data. Accordingly, the main text was revised to emphasize that all MRI values for the AD subtype cluster were lower than the MRI ROI values of the remaining dataset.

