## Supplemental file for "Alzheimer’s subtypes A supervised, unsupervised, multimodal, multilayered embedded recursive (SUMMER) AI study"

| Table 1. ROI intensity measurements | | |
|---|---|---|
| | **Two ROIs histogram measurements (Between the hemispheres, left and right)** | **One ROI histogram** |
| 1. Statistical measurements | | Min, max, std, median, top 25th percentile, top 20th percentile, top 10th percentile, bottom 10th percentile, bottom 20th percentile, and bottom 25th percentile. |
| 2. Euclidean Distance = Calculating Distances Between 2 Histograms by Using Euclidean Distance. | √ | |
| 3. The entropy of a histogram measures the amount of uncertainty, randomness, or information content within the distribution of the histogram. It is a concept from information theory that quantifies how spread out or unpredictable the data is. | | √ |
| 4. Difference between two entropies of histograms<br>(Jensen-Shannon divergence) | √ | |
| 5. Percentage of mutual area (vs. both histograms) | √ | |
| 6. Percentage of mutual areas (vs. the right) | √ | |
| 7. Percentage of mutual areas (vs. the left) | √ | |

| Table 2. Selected ROIs unsupervised analysis | | | | | |
|---|---|---|---|---|---|
| **ROI** | **Mean CN** | **Mean AD** | **SD CN** | **SD AD** | **P value** |
| **Men group** | | | | | |
| Top 25 Supracalcarine Cortex | 2.644 | 2.144 | 0.308 | 0.18 | 1.67E-16 |
| SD Occipital_Fusiform | 0.727 | 0.535 | 0.134 | 0.066 | 1.43E-15 |
| Top 10 Lingual Gyrus | 2.866 | 2.360 | 0.370 | 0.170 | 2.80E-15 |
| Top 20 Parietal Operculum | 2.599 | 2.235 | 0.197 | 0.167 | 2.22E-14 |
| SD Right Hippocampus | 0.780 | 0.594 | 0.131 | 0.088 | 1.86E-12 |
| Bottom 10 Frontal Operculum | 0.213 | 0.249 | 0.158 | 0.143 | 0.2870 |
| Bottom 10 Right Amygdala | 0.267 | 0.294 | 0.285 | 0.156 | 0.55 |
| **Wome group** | | | | | |
| Top 25 Parahippocampal Gyrus | 2.122 | 1.584 | 0.345 | 0.093 | 5.84E-12 |
| Entropy Parahippocampal Gyrus | 3.498 | 3.929 | 0.318 | 0.110 | 9.19E-10 |
| Top 20 Temporal Fusiform | 2.242 | 1.704 | 0.467 | 0.142 | 6.23E-08 |
| Bottom 20 Heschl Gyrus | 0.392 | 0.197 | 0.174 | 0.077 | 2.90E-07 |
| Mean Right Caudate | 1.567 | 1.419 | 0.138 | 0.078 | 1.51E-06 |
| Entropy Right Hippocampus | 3.768 | 3.994 | 0.229 | 0.077 | 2.37E-06 |
| Entropy Right Caudate | 3.717 | 3.840 | 0.119 | 0.077 | 4.89E-06 |
| Median Right Thalamus | 1.995 | 1.750 | 0.266 | 0.140 | 1.94E-05 |
| Entropy Superior Temporal | 3.949 | 4.037 | 0.088 | 0.077 | 3.29E-05 |
| Note: CN- Cognitve normal, AD – Alzheimer's disease, ROI – Region of interest, Top-top 25$^{th}$ percentile, Top-top 20$^{th}$ percentile, Top-top 10$^{th}$ percentile, Bottom-bottom 10$^{th}$ percentile. SD – Standard deviation | | | | | |

| Table 3. Unsupervised analysis, percent of the participants | | | |
|---|---|---|---|
| | Rest of the data | Cluster | Percent of the AD in the cluster |
| Men | 75% | 25% | 92.3% |
| Women | 61.85% | 38.15% | 86.2% |

### Unsupervised machine learning applications (Clusters)

### Women group

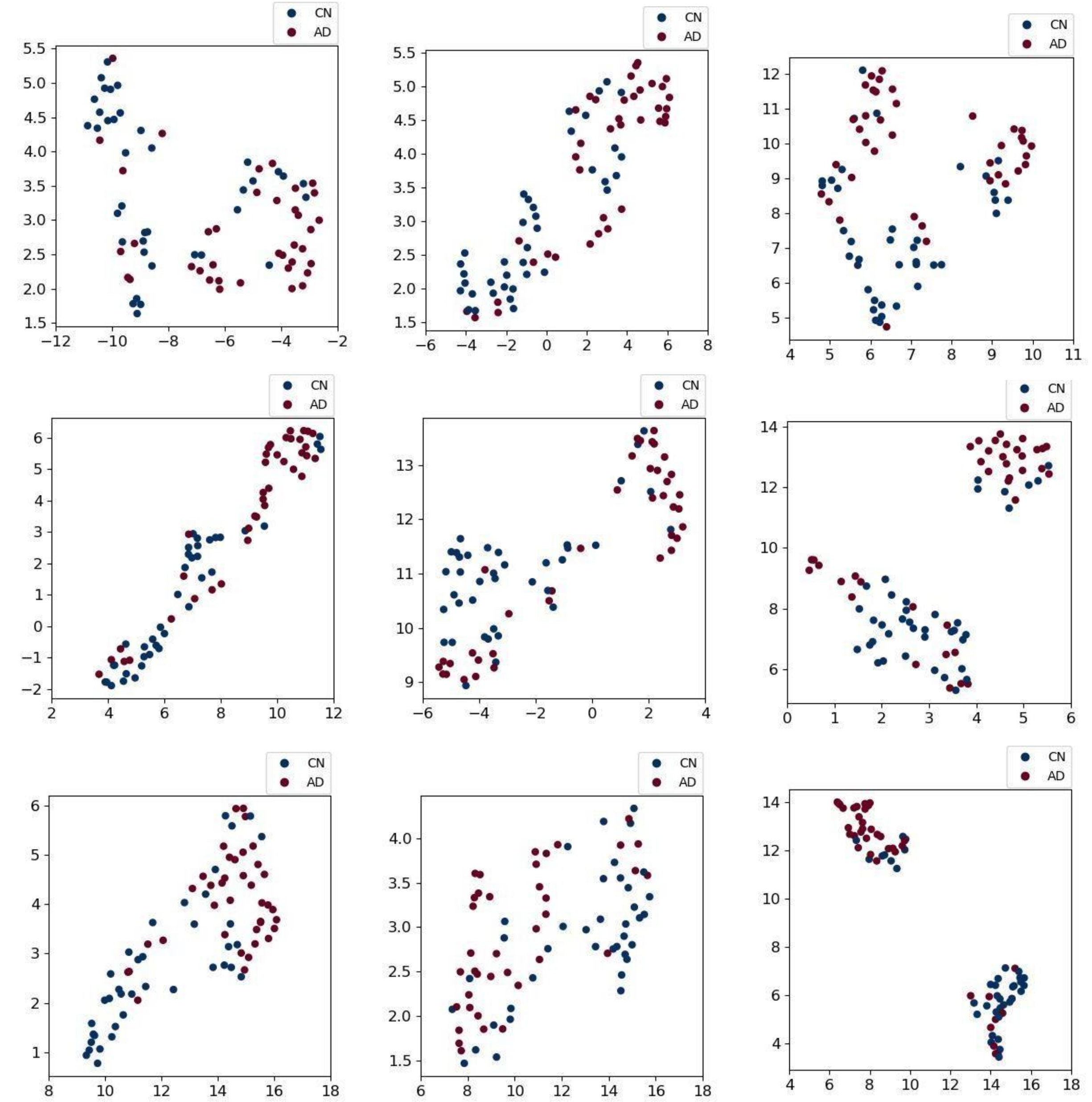

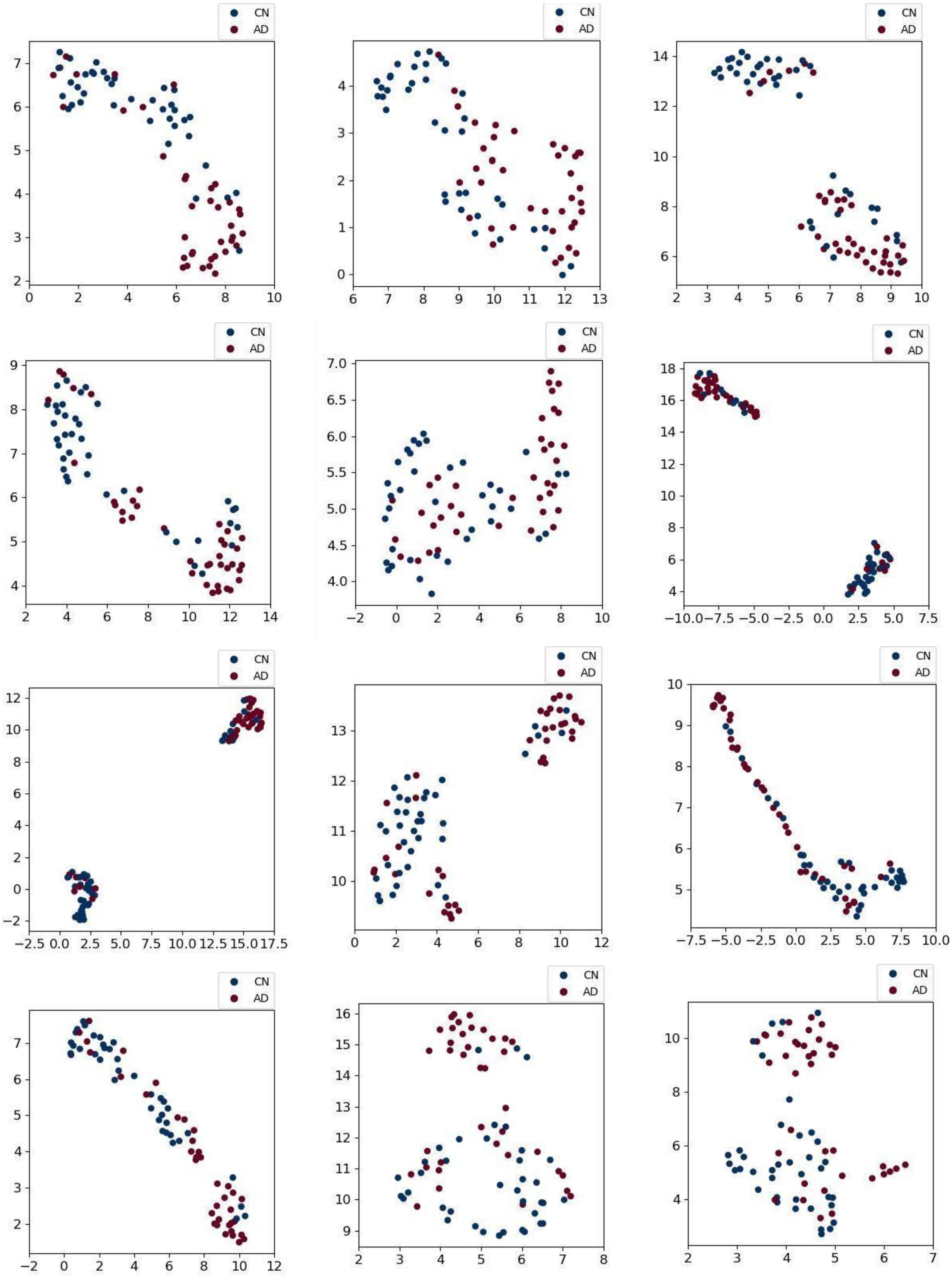
CN
AD

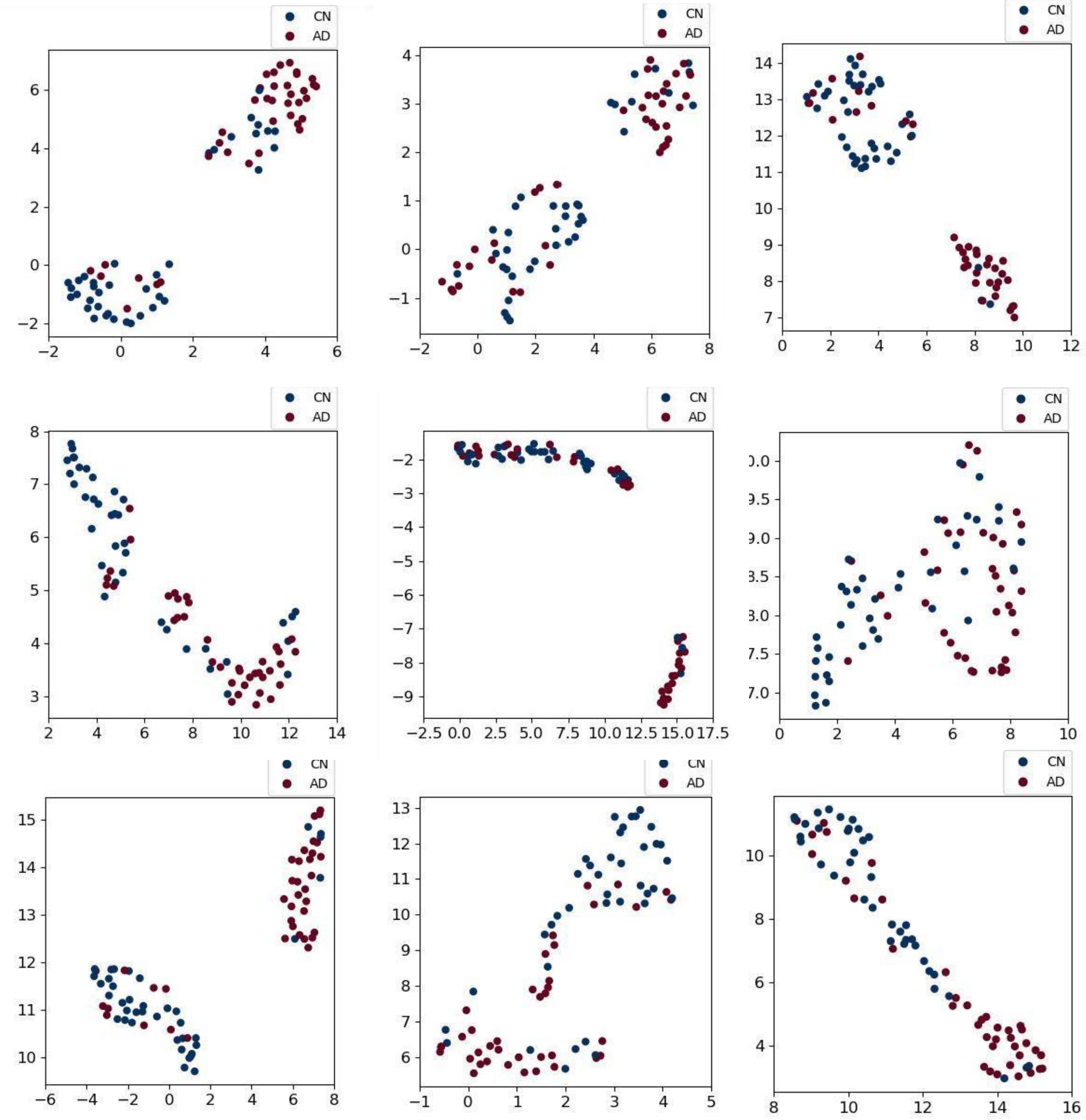
CN
AD

### Men group

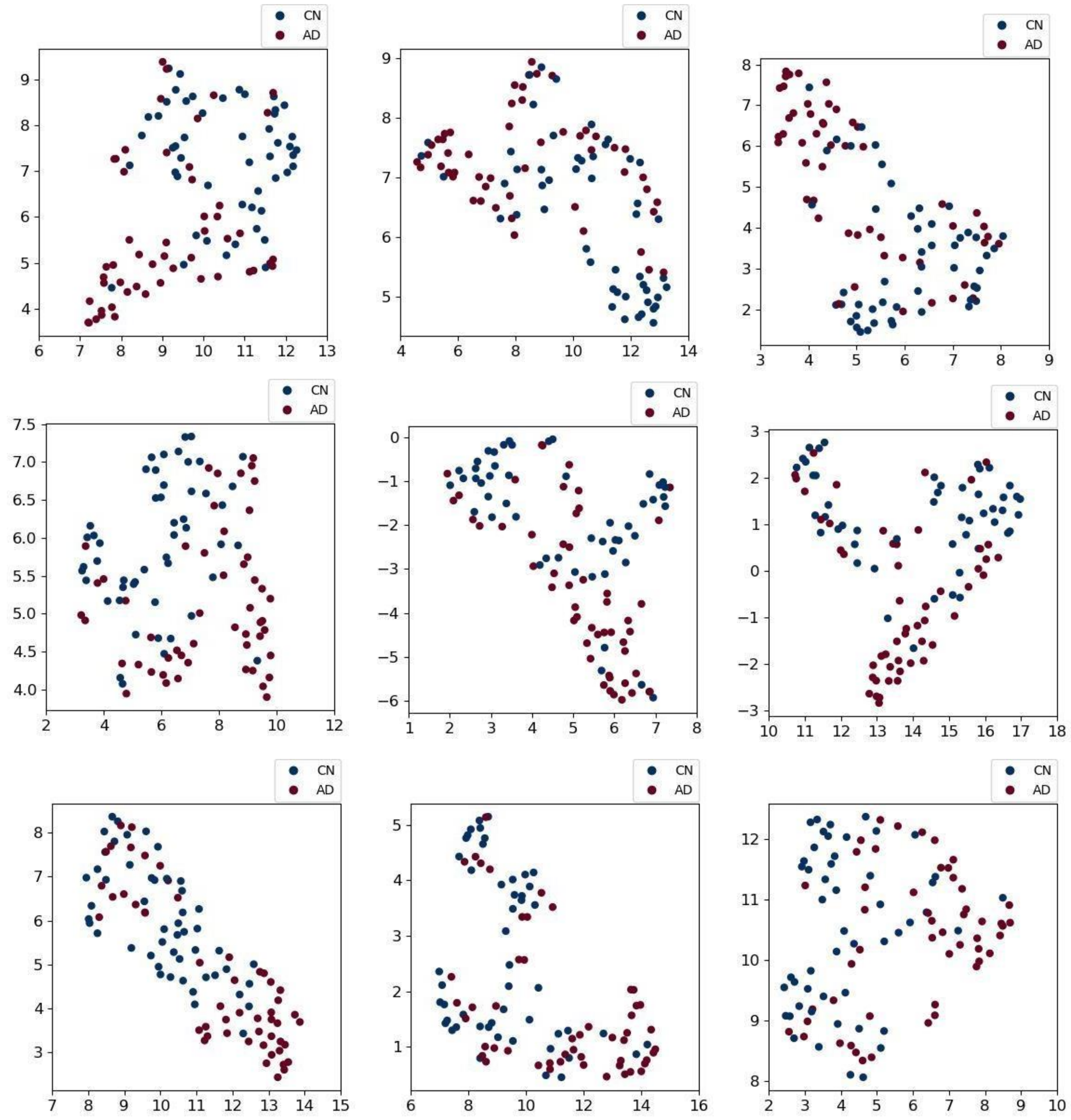

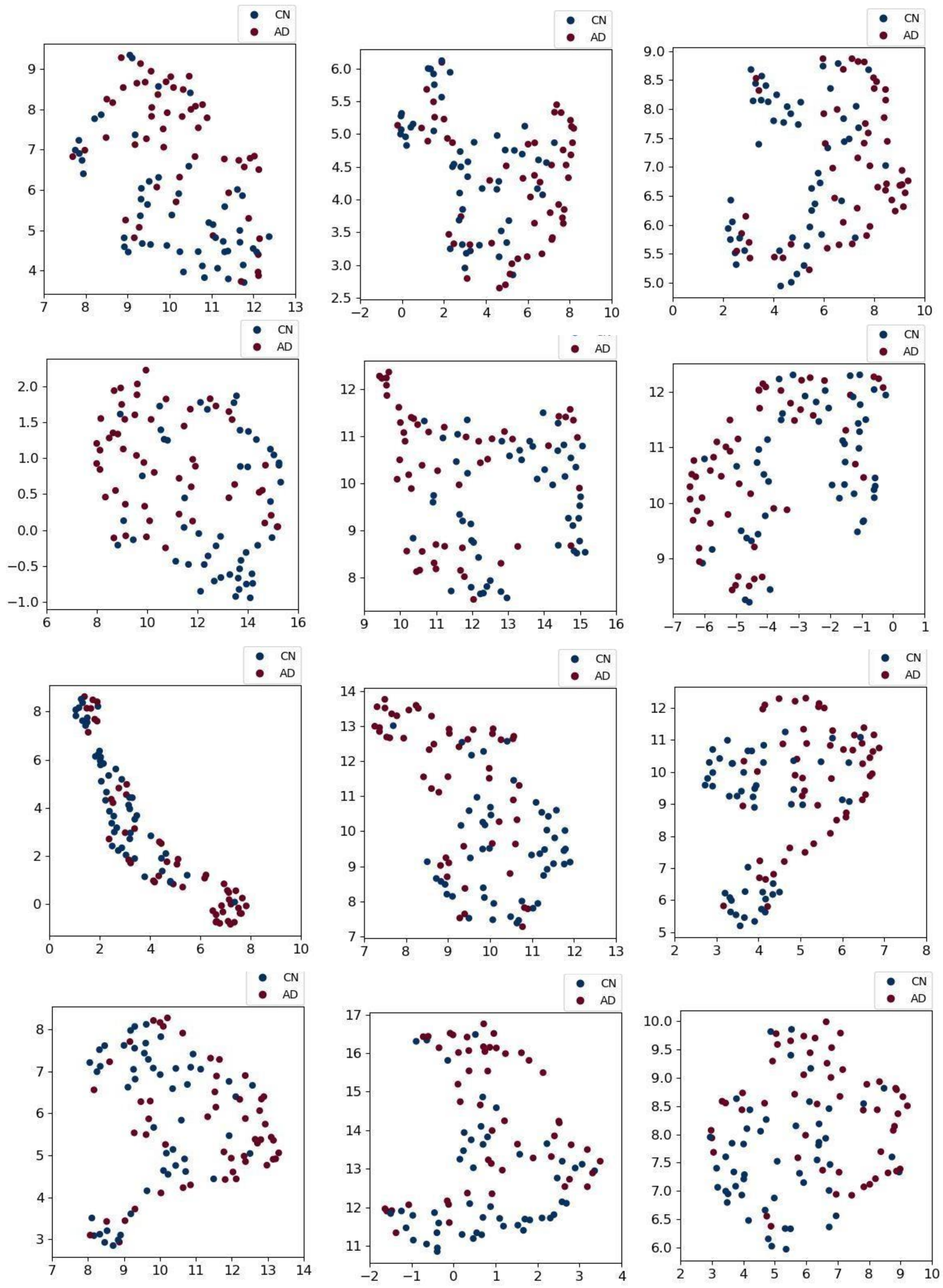
CN
AD

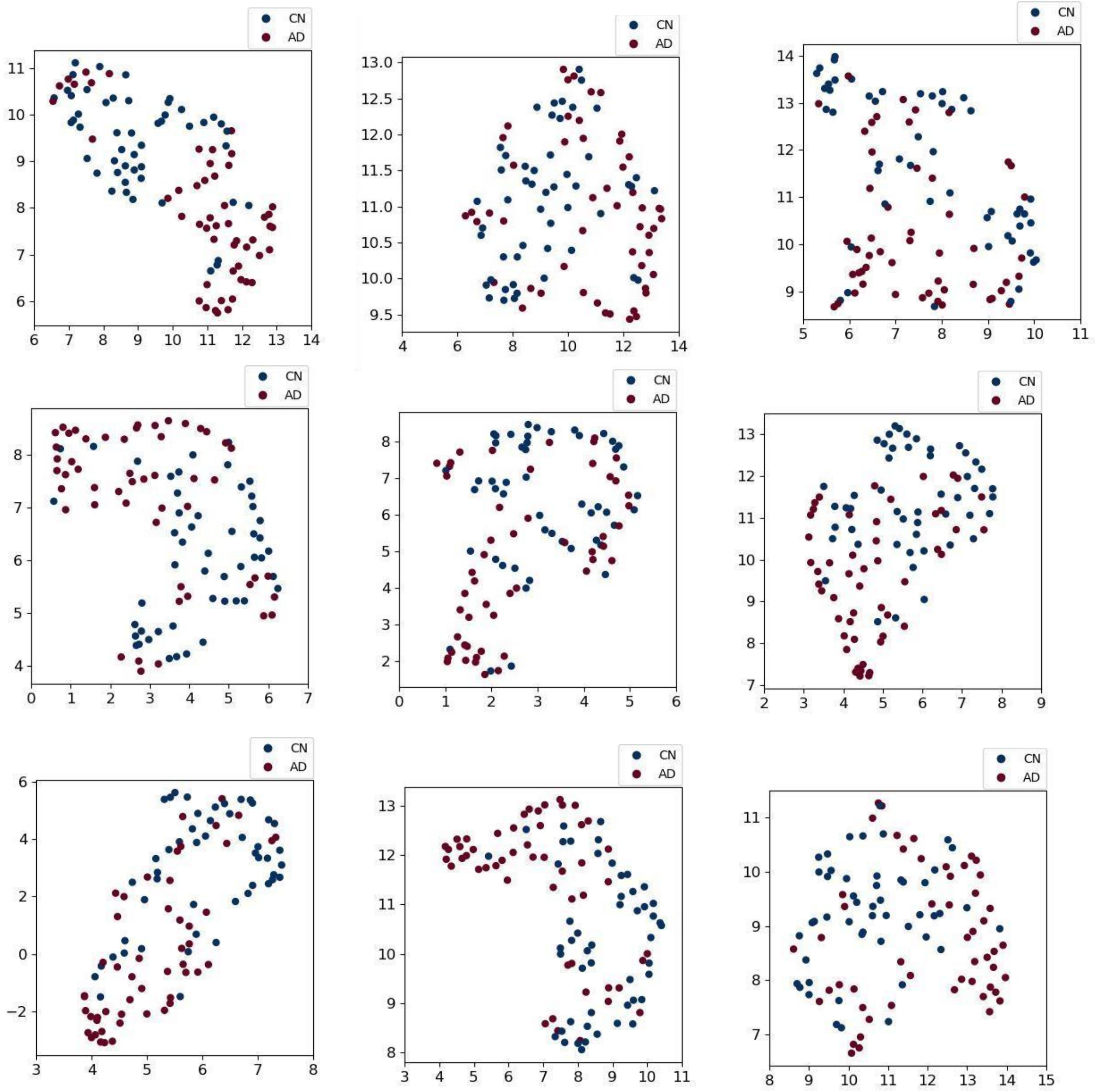
CN
AD

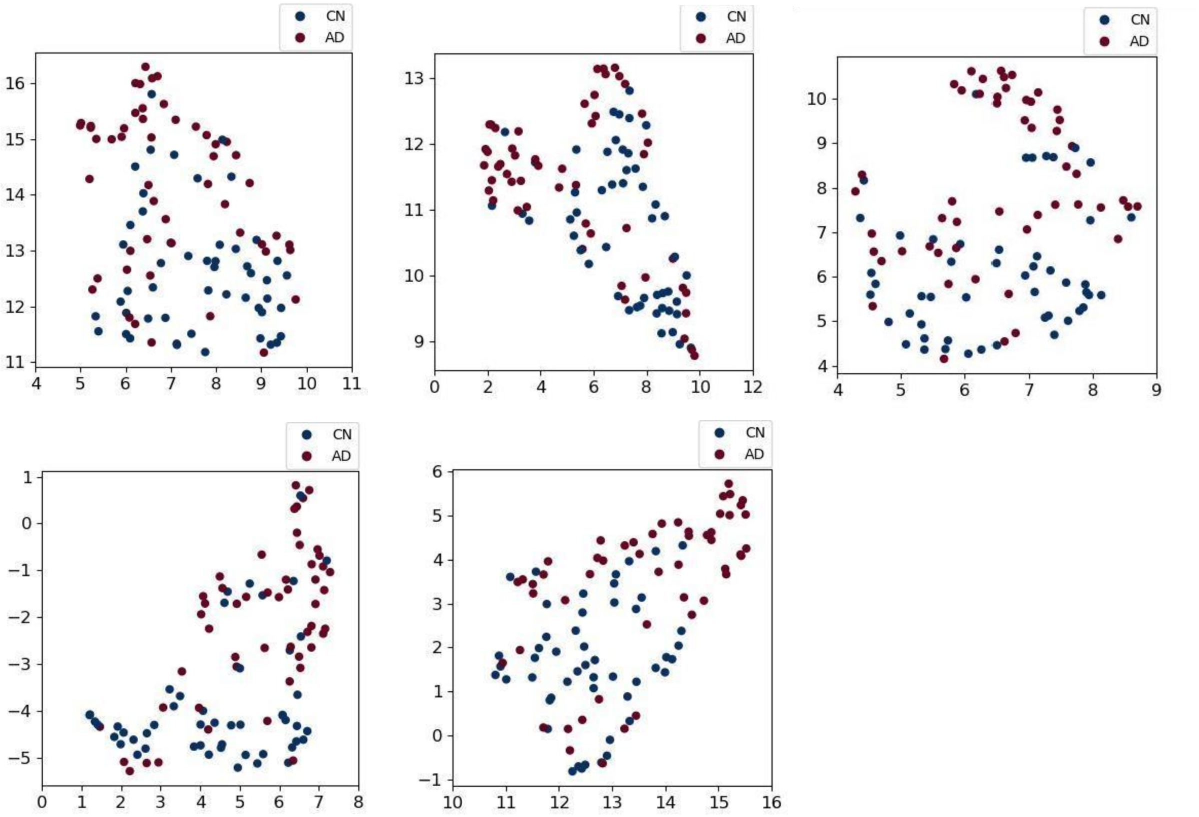
CN
AD

| Table 4: Supervise analysis, selected SNPs | | | | | | | | |
|---|---|---|---|---|---|---|---|---|
| Weight ranking | ID | Cromozone | Position | Ref | Alt column only | Gene | MAF: | Highest population MAF: |
| | **women** | | | | | | | |
| 1 | rs11689050 | 2 | 19472420 | G | A | LINC01808 | 0.07 | 0.23 |
| 2 | rs1874161 | 5 | 158188702 | C | A | | 0.5 | 0.11 |
| 3 | rs10216685 | 8 | 76507533 | G | A,T | | 0.14 | 0.46 |
| 4 | rs6775796 | 3 | 74654538 | G | A | | 0.21 | 0.42 |
| 5 | rs7306261 | 12 | 87879250 | T | A,C,G | | 0.26 | 0.43 |
| 6 | rs2139011 | 2 | 122930569 | A | G,T | | 0.11 | 0.22 |
| 7 | rs10069215 | 5 | 111050943 | G | A,T | | 0.13 | 0.23 |
| 8 | rs2422973 | 20 | 4782055 | A | C,G | RASSF2 | 0.18 | 0.35 |
| 9 | rs17456902 | 10 | 68124249 | G | A,T | | 0.10 | 0.24 |
| 10 | rs755982 | 8 | 117561710 | C | T | | 0.09 | 0.2 |
| | **Men** | | | | | | | |
| 1 | rs7722673 | 5 | 168417231 | G | T | WWC1 | 0.05 | 0.12 |
| 2 | rs1538608 | 10 | 9889061 | A | G | | 0.13 | 0.35 |
| 3 | rs2396274 | 2 | 226025312 | A | C,G,T | | 0.16 | 0.41 |
| 4 | rs1122269 | 20 | 61668780 | C | A,G,T | CDH4 | 0.18 | 0.48 |
| 5 | rs17131515 | 1 | 91650462 | C | A,T | | | 0.49 |
| 6 | rs10050582 | 5 | 63705116 | C | A,T | | 0.06 | 0.2 |
| 7 | rs17513184 | 5 | 54290040 | C | T | | 0.03 | 0.12 |
| 8 | rs12130821 | 1 | 163229800 | C | A,T | RGS5 | | 0.22 |
| 9 | rs9425322 | 1 | 183864927 | G | A,T | RGL1 | | 0.24 |
| 10 | rs17794576 | 21 | 21133878 | G | A,C,T | NCAM2 | 0.12 | 0.32 |

| Table 5. Unseen ADNI data, Supervised Genetics Model results | | | |
|---|---|---|---|
| Subject Category | Corrected Predicted (True Positive) | Not corrected predicted (False Negative) | Overall |
| AD men | 45 | 33 | 78 |
| CN men | 188 | 99 | 287 |
| AD women | 61 | 11 | 72 |
| CN women | 227 | 139 | 366 |
